## Supplementary Info for "Interaction of Genetic Variants Activates Latent Metabolic Pathways in Yeast"

**This supplementary information file includes**

Supplementary Methods and Results

Supplementary Data 1 and 2

Legends for Supplementary Data 3-10

Supplementary Figures

**SUPPLEMENTARY METHODS**

**RNA Extraction Protocol**

RNA was extracted from yeast cells using Qiagen RNeasy mini kit (Cat No.74106). The cell pellet was resuspended in buffer RLT and incubated at room temperature for 5 min to achieve complete lysis. The lysate mixed with half volume of absolute alcohol was loaded into the RNeasy spin column placed in a 2 ml collection tube. The tubes were centrifuged at 8,000 rpm for 1 min, and flow through discarded. On column DNase I (Cat No.79254), treatment and subsequent column washes were performed according to manufacturer’s protocol. RNA was eluted from the column using Nuclease free water.

The concentration and purity of RNA quantified using Nanodrop Spectrophotometer (Thermo Scientific; 2000). The integrity of RNA in the samples assessed on Tapestation (Agilent). RNA concentration quantified using Qubit RNA HS assay kit (Q32855).

**Library Preparation and Sequencing**

RNA sequencing libraries were prepared with Illumina-compatible NEBNext® Ultra™ II Directional RNA Library Prep Kit (New England BioLabs, MA, USA) at Genotypic Technology Pvt. Ltd., Bangalore, India.

100- 500 ng of total RNA was taken for mRNA isolation, fragmentation and priming. Fragmented and primed mRNA was further subjected to first-strand synthesis followed by second-strand synthesis. The double-stranded cDNA was purified using NEBNext sample purification beads. Purified cDNA was end-repaired, adenylated and ligated to Illumina adapters as per NEBNext® Ultra™ II Directional RNA Library Prep protocol followed by second strand excision using USER enzyme at 37 ˚C for 15mins.

Illumina Universal Adapters used in the study were:

5’-AATGATACGGCGACCACCGAGATCTACACTCTTTCCCTACACGACGCTCTTCCGATCT-3’ and

Index Adapter:

5’-GATCGGAAGAGCACACGTCTGAACTCCAGTCAC [INDEX] ATCTCGTATGCCGTCTTCTGCTTG-3’.

[INDEX] – Unique sequence to identify sample-specific sequencing data.

Adapter ligated cDNA was purified using NEBNext beads and was subjected to 11 cycles for Indexing-(98˚C for 30 sec, cycling (98˚C for 10sec, 65˚C for 75sec) and 65˚C for 5min) and enrich the adapter-ligated fragments. Final PCR products (sequencing library) were purified with NEBNext beads, followed by library quality control check. Illumina-compatible sequencing libraries were quantified by Qubit fluorometer (Thermo Fisher Scientific, MA, USA) and fragment size distribution was analysed on Agilent 2200 TapeStation.

**Proteomic sample processing**

Cell pellets of *S. cerevisiae* were lysed in 6 M guanidinium·HCl, 5 M tris(2-carboxyethyl) phosphine, 10 M chloroacetamide, and 100 M Tris·HCl (pH = 8.5), disrupted mechanically and heated to 99°C. After centrifugation, the cell-free lysates were diluted with 50 M ammonium bicarbonate and subjected to a bicinchoninic acid (BCA) assay to estimate protein concentrations. Trypsin and LysC digestion mix (Promega) was added to 20 μg protein of each sample and incubated for 8 hours. Trifluoroacetic acid was added to halt digestion, and the samples were desalted using C18 resin (Empore, 3M) before HPLC-MS analysis. HPLC-MS analysis of the samples was performed on an Orbitrap Exploris 480 instrument (Thermo Fisher Scientific), preceded by an EASY-nLC 1200 HPLC system (Thermo Fisher Scientific). For each sample, 0.5 μg of peptides were captured on a 2 cm C18 trap column (Thermo Fisher 164946). Subsequently, separation was executed using a 70 min gradient from 8% (v/v) to 48% (v/v) of acetonitrile in 0.1% (v/v) formic acid on a 15 cm C18 reverse-phase analytical column (Thermo EasySpray ES904) at a flow rate of 250 nL/min. For data-independent acquisition, the mass spectrometer was run with the HRMS1 method as previously described^1^, preceded by the FAIMS Pro Interface (Thermo Fisher Scientific) with a compensation voltage (CV) of -45 V, and any modifications are mentioned below. Full MS1 spectra were collected at a resolution of 120,000 and a scan range of 400-1,000 m/z, with the maximum injection time set to auto. MS2 spectra were obtained at a resolution of 60,000, with the maximum injection time set to auto and the collision energy set to 32. Each cycle consisted of three DIA experiments, each covering a range of 200 m/z with a window size of 6 m/z and a 1 m/z overlap, while a full MS scan was obtained between experiments.

**Metabolomic sample processing and analysis:**

For LC-MS/MS analysis, dried extracts were reconstituted in 200 µL of LC-MS-grade water containing 0.1% formic acid. Samples were vortexed for 30 seconds, spun briefly, and sonicated for 2 minutes. After centrifugation at 14,000 × g for 10 minutes at 4 °C, the supernatant was transferred to a fresh tube and diluted 5-fold with 0.1% formic acid. A 10 µL aliquot was injected into the LC-MS system.

Chromatographic separation was performed using an Agilent 1290 Infinity II UHPLC system equipped with a C18 column (4 µm, 4.6 × 150 mm). The mobile phases consisted of water with 0.1% formic acid (A) and methanol with 0.1% formic acid (B) at a 0.5 ml/min flow rate. The column oven and autosampler were maintained at 45 °C and 4 °C, respectively. The following gradient was applied: 0–3 min, 2% B; 3–12 min, linear increase to 35% B; 12–15 min, linear increase to 90% B; 15–16 min, 90% B; 16–16.1 min, decrease to 5% B; 16.1–20 min, re-equilibration at 5% B.

Detection was performed on an Agilent 6495 Triple Quadrupole mass spectrometer operating in positive ionization mode using multiple reaction monitoring (MRM). Data acquisition and quantification were conducted using Agilent MassHunter Workstation Quantitative Analysis software (version 10.1). To monitor instrument performance and ensure data quality, three pooled quality control (QC) samples—prepared by combining equal aliquots from all experimental samples—were analyzed at the beginning, middle, and end of the run. The coefficient of variation (CV) for each amino acid was calculated, and it was ensured that the CV remained below 15% for all analytes (Supplementary Data 9).

**Integrating protein expression data into yeast genome-scale metabolic model**

The protein constraint metabolic models for each strain and their respective time phases were generated by integrating the protein expression data into the latest genome-scale metabolic model of yeast (yeast9)^2^ (<https://github.com/SysBioChalmers/yeast-GEM>) using the model extraction method iMAT^3^ using lower threshold as the minimum value of the expression array and upper threshold as the minimum value of top 10 % of reaction’s expression array for that particular strain and time point. The bounds of exchange reactions of the GEM were changed to reflect the early stages of the sporulation state appropriately, as followed in our previous study^4^. For this, we restricted the intake of glucose and nitrogen (Lower bound = Upper bound = 0 [mmol/ (g DW h)]) while allowing an unrestricted supply of oxygen and acetate (Lower bound = -1000 [mmol/ (g DW h)]). The metabolic heterogeneity between the context-specific models was studied by calculating the Jaccard index between the models.

**Genome-scale differential flux analysis to predict upregulated and downregulated reactions**

The optGpSampler^5^ in COBRApy version 0.10^6^ was used to sample 10000 flux solutions with a thinning factor set to 100 on all the generated SNP and time-specific metabolic models.

Let $X_{SNP}$ and $X_{SS}$ be the flux distributions of a particular reaction in SNP-specific (MM, TT, and MMTT) and SS models for each time point (0h and 2h30m). The flux change was calculated for each reaction as follows:

$$Flux change= \frac{\bar{X}_{SNP}- \bar{X}_{SS}}{\left| \bar{X}_{SNP}+\bar{X}_{SS} \right|}$$

where, $\bar{X}_{SNP}$and $\bar{X}_{ss}$, the arithmetic means of the flux distributions for a given reaction in SNP and null model, respectively.

To identify significantly altered reactions in each SNP-specific model compared to the null model, we employed a two-sided Kolmogorov-Smirnov test with a significance threshold 0.05, adjusted for multiple comparisons using the Benjamini-Hochberg method. From the set of dysregulated reactions, we classified those with a flux change greater than 0.82 as upregulated and those with a flux change below -0.82 as downregulated^4,7^, corresponding to a tenfold increase or decrease in flux in the SNP model relative to the null model. The outline of steps involved in generating context-specific models and GS-DFA analysis is shown in Supplementary Fig. 18A.

**SUPPLEMENTARY RESULTS**

We constructed context-specific metabolic models for the SS, MM, TT, and MMTT yeast strains at two-time points: 0 hours and 2 hours 30 minutes. This was achieved by integrating SNP-specific protein abundance data into the Yeast GEM (Yeast9) using iMAT (Integrative Metabolic Analysis Tool). iMAT categorises protein expression levels into high, moderate, and low, creating a subnetwork enriched with reactions driven by highly expressed proteins while minimising the inclusion of reactions with low expression. It aims to maximise the consistency between reaction activity and protein expression, assigning non-zero flux to active reactions and zero to inactive ones. The SNP-specific models generated by iMAT, including the number of genes, reactions, and metabolites in each, are summarised in Supplementary Fig. 18B.

We calculated the Jaccard similarity index to evaluate the variability among these models. We observed that the models at 0 hours (SS-0h, MM-0h, TT-0h, and MMTT-0h) and the 2h30m model of the low-sporulating strain (SS-2h30m) are highly similar. In contrast, the models for high-sporulating strains at 2h30m (MM-2h30m, TT-2h30m, and MMTT-2h30m) exhibited high similarity (Supplementary Fig. 18C).

We then focused on identifying reactions upregulated in the MM, TT, and MMTT strains compared to the SS strain at both time points (Supplementary Data 10). Specifically, we investigated the flux through the arginine biosynthesis pathway, as we had previously shown that the MMTT strain exhibits unique regulation of this pathway, with increasing allocation during sporulation. Additionally, we aimed to identify pathways uniquely enriched in the MMTT strain, potentially due to genetic interactions between the *MKT1*^89G^ and *TAO3*^4477C^ SNPs.

Our analysis revealed that intracellular flux through the arginine biosynthesis pathway, particularly Argininosuccinate lyase (*ARG4*), is inactive at 0 hours in all strains but becomes active in the MM-2h30, TT-2h30, and MMTT-2h30 models while remaining inactive in the SS-2h30m model. This suggests that *ARG4* flux may be critical for efficient sporulation (Supplementary Fig. 19A, Supplementary Fig. 19B). Furthermore, flux enrichment analysis of upregulated reactions identified using GS-DFA revealed that the steroid biosynthesis pathway is uniquely enriched in the MMTT strain (adjusted p-value = 7.59E-08). The intracellular flux distribution of key reactions in the steroid biosynthesis pathway is illustrated in Supplementary Fig. 20.

**SUPPLEMENTARY DATA**

**Supplementary data 1:** List of strains used in the study

| Strain identifier | Strain | Parent strain | Reference |
| --- | --- | --- | --- |
|  | S288c |  | Deutschbauer and Davis^8^ |
|  | SK1 |  | Deutschbauer and Davis ^8^ |
| SHS306 | MM (*MKT1*-89G (a/alpha)) | SHS304 x SHS305 (M strain) | Gupta *et al*. ^9^ |
| SHS327 | TT (*TAO3*-4477C (a/alpha)) | SHS325 x SHS326 (clean BC3T strain) | Gupta *et al.*^10^ |
| SHS884 | MmTt strain [*MKT1*(89G)/*MKT1*(89A) *TAO3*(4477C)/*TAO3*(4477G)] | by mating M strain and T strain | This study |
| SHS885 | MT strain (a) | by sporulating MmTt strain | This study |
| SHS886 | MT strain (alpha) | by sporulating MmTt strain | This study |
| SHS887 | MMTT strain (diploid) | by crossing SHS885 and SHS886 | This study |
| SHS888 | S288c, ∆*arg4*::HygMX4 (a) | S288c (a) | This study |
| SHS889 | M strain , ∆*arg4*::HygMX4 (alpha) | M strain (alpha) | This study |
| SHS890 | T strain , ∆*arg4*::HygMX4 (a) | T strain (a) | This study |
| SHS891 | MT strain, ∆*arg4*::HygMX4 (a) | MT strain (a) | This study |
| SHS892 | S288c, ∆*arg5*6::HygMX4 (a) | S288c (a) | This study |
| SHS893 | M strain , ∆*arg5*6::HygMX4 (alpha) | M strain (alpha) | This study |
| SHS894 | T strain , ∆*arg5*6::HygMX4 (a) | T strain (a) | This study |
| SHS895 | MT strain, ∆*arg5*6::HygMX4 (a) | MT strain (a) | This study |
| SHS896 | S288c, ∆*arg4*::HygMX4 (Diploid) | S288c (a) | This study |
| SHS897 | MM strain , ∆*arg4*::HygMX4 (Diploid) | M strain (alpha) | This study |
| SHS898 | TT strain , ∆*arg4*::HygMX4 (Diploid) | T strain (a) | This study |
| SHS899 | MMTT strain, ∆*arg4*::HygMX4 (Diploid) | MT strain (a) | This study |
| SHS900 | S288c, ∆*arg5*6::HygMX4 ((Diploid) | S288c (a) | This study |
| SHS901 | MM strain , ∆*arg5*6::HygMX4 (Diploid) | M strain (alpha) | This study |
| SHS902 | TT strain , ∆*arg5*6::HygMX4 (Diploid) | T strain (a) | This study |
| SHS903 | MMTT strain, ∆*arg5*6::HygMX4 (Diploid) | MT strain (a) | This study |

**Supplementary data 2:** List of primers used in the study

| Primer name | Primer sequence 5'-3' | Description |
| --- | --- | --- |
| HS3501 | GCTCAAAAGC AGGTAACTAT ATAACAAGAC TAAGGCAAAC cagctgaagcttcgtacgct | FP *ARG4* deletion |
| HS3502 | CCAGACCTGATGAAATTCTTGCGCATAACGTCGCCATCTG cataggccactagtggatctg | RP *ARG4* deletion |
| HS3503 | ACGTATTTTAGCCTGGTTGT | FP with BN72 (Junction amplification *ARG4*) |
| HS3504 | ATCCTGCAGATTGGGTCACA | RP with JM37 (Junction amplification *ARG4*) |
| HS3505 | AGAAAGAGAG ATCTGAAGTG AGAAATAAGT CTGCATCATT cagctgaagcttcgtacgct | FP *ARG5*6 deletion |
| HS3506 | GTCTCATGTGACTGAGCTGCAGATCTACTATACACAGTGA cataggccactagtggatctg | RP *ARG5*6 deletion |
| HS3507 | GGAAAGGTATCTGGGAAAGG | FP with BN72 (Junction amplification *ARG5*6) |
| HS3508 | AAACGCGTAAACAAGCTCGT | RP with JM37 (Junction amplification *ARG5*6) |
| HS1013 | CTGAATAATTGTACCCTGGA | Fwd primer to check *MKT1* 89A SNP with HS1015 |
| HS1014 | CTGAATAATTGTACCCTGGG | Fwd primer to check *MKT1* 89G SNP with HS1015 |
| HS1015 | GTTGAAACCAAGAGGAGTAA | Common reverse primer with HS1013 and HS1014 |
| HS1016 | TGATCTACTTTCAGCTGTTG | Forward primer to check *TAO3* 4477G SNP with HS1018 |
| HS1017 | TGATCTACTTTCAGCTGTTC | Forward primer to check *TAO3* 4477C SNP with HS1018 |
| HS1018 | GCTAAAGGAACCATGTATTT | Common reverse primer with HS1016 and HS1017 |
| HS1127 | CTCCACTTCAAGTAAGAGTTTGGGT | forward primers specific to the MATa |
| HS1128 | TTACTCACAGTTTGGCTCCGGTGT | forward primers specific to the MATalpha |
| HS1129 | GAACCGCATGGGCAGTTTACCTTT | common reverse primer to MAT alleles |

**Supplementary Data 3:** Differentially expressed genes identified using DESeq2.

**Supplementary Data 4:** Differentially expressed genes identified using the DESeq2 LRT method.

**Supplementary Data 5:** List of genes present in each cluster for SS and MMTT strain data identified using DPGP clustering.

**Supplementary Data 6:** Differentially expressed proteins MM, TT and MMTT strain at 0 h identified in comparison to the SS strain.

**Supplementary Data 7:** Differentially expressed proteins in MM, TT and MMTT strains at 2 h 30 min identified in comparison to the SS strain

**Supplementary Data 8:** Differentially expressed proteins in SS, MM, TT and MMTT strains at 2 h 30 min identified compared to the 0th hour of each strain, respectively.

**Supplementary Data 9:** Raw intensity data of intracellular amino acids obtained for SS, MM, TT and MMTT strains at three timepoints (0 h, 2 h 30 min, 8 h) during sporulation.

**Supplementary Data 10:** List of upregulated reactions identified using GS-DFA analysis.

**SUPPLEMENTARY FIGURES**

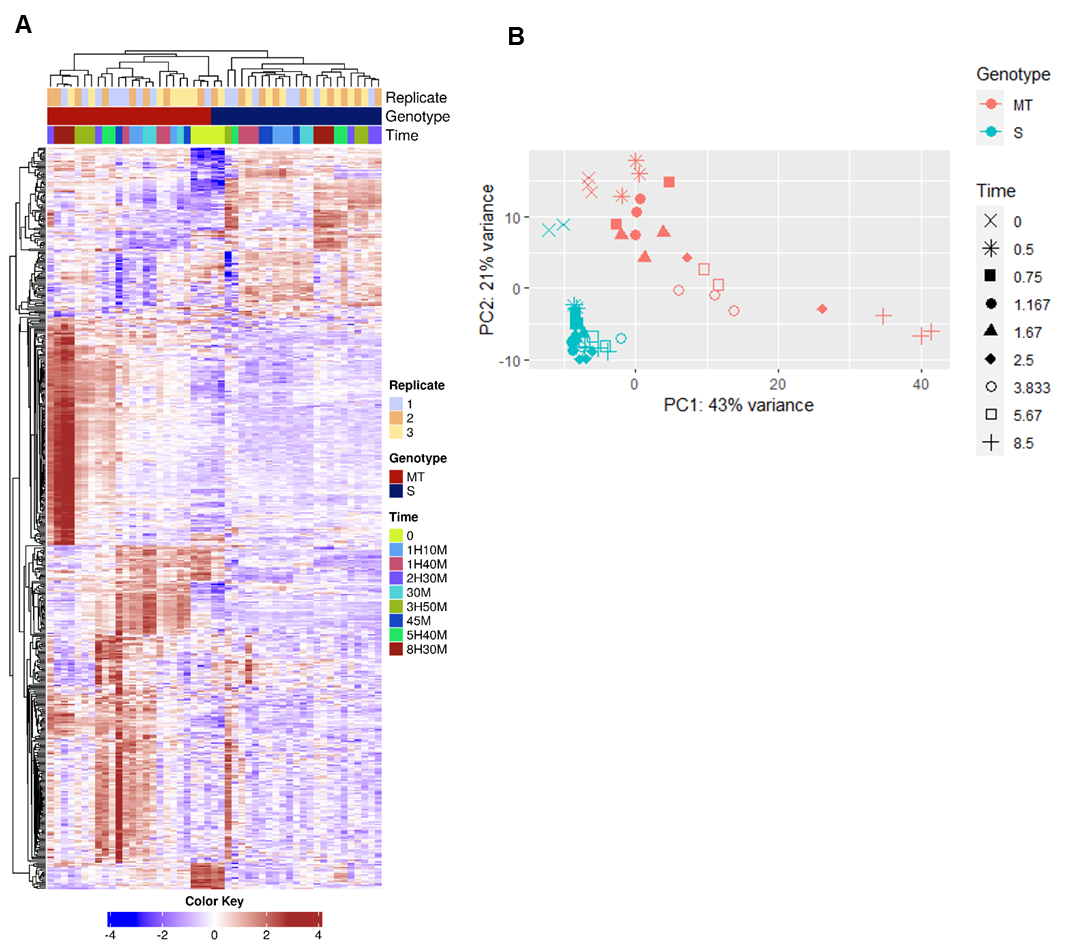

**Supplementary Fig. 1:** **Distinct temporal transcriptional profiles in SS and MMTT strains during sporulation.** (A) Heatmap representing the hierarchically clustered gene expression levels of SS and MMTT strains across 9 time points for the 300 most variable genes. (B) PCA plot based on the 300 most variable genes.

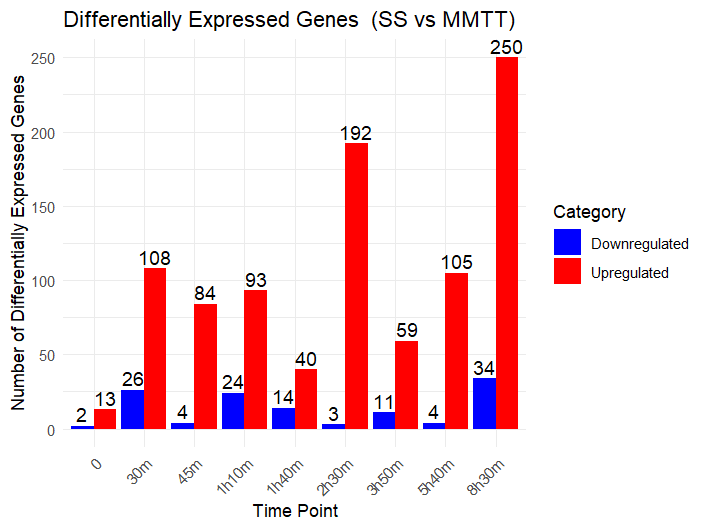

**Supplementary Fig. 2:** The number of differentially expressed genes in the MMTT strain when compared to the SS strain at each time point during sporulation.

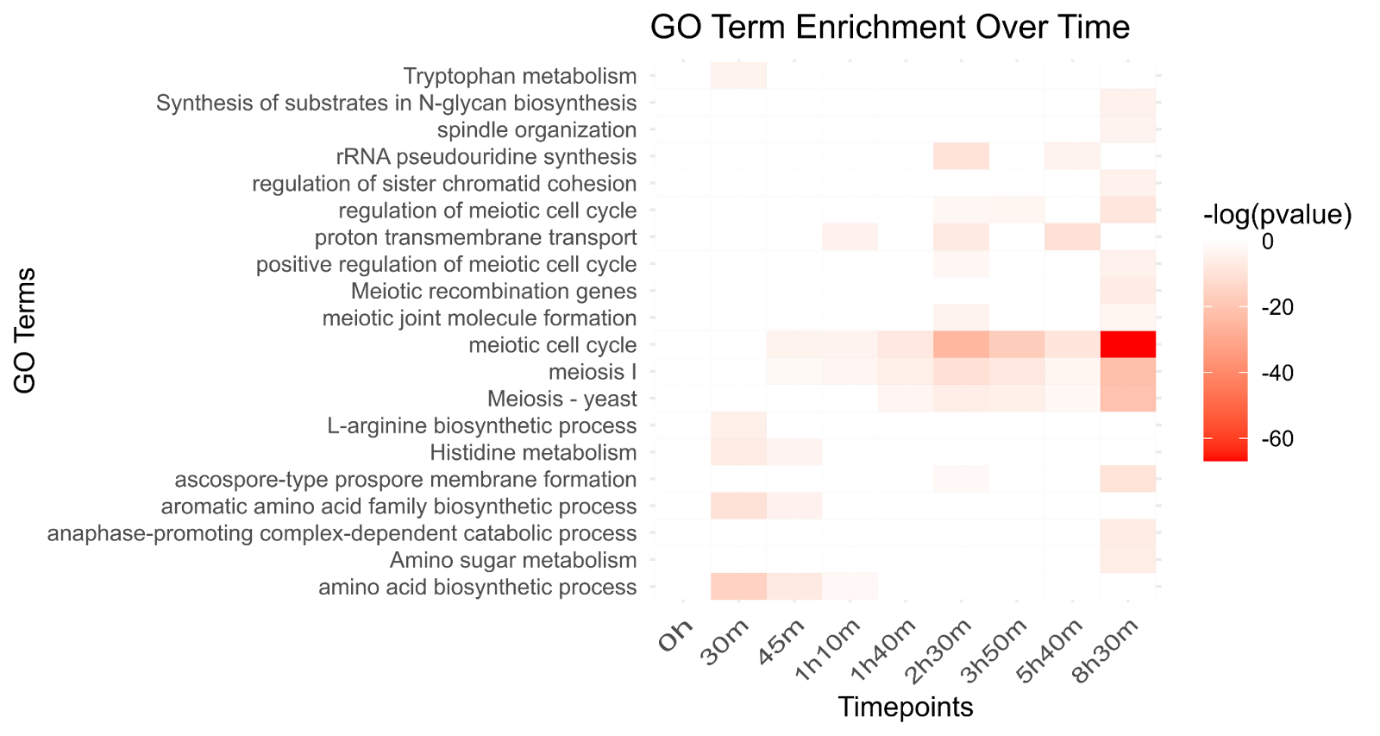

**Supplementary Fig. 3:** GO enrichment analysis for upregulated genes in the MMTT strain across each time point in comparison with the SS strain. The heatmap represents the p-value of each GO term.

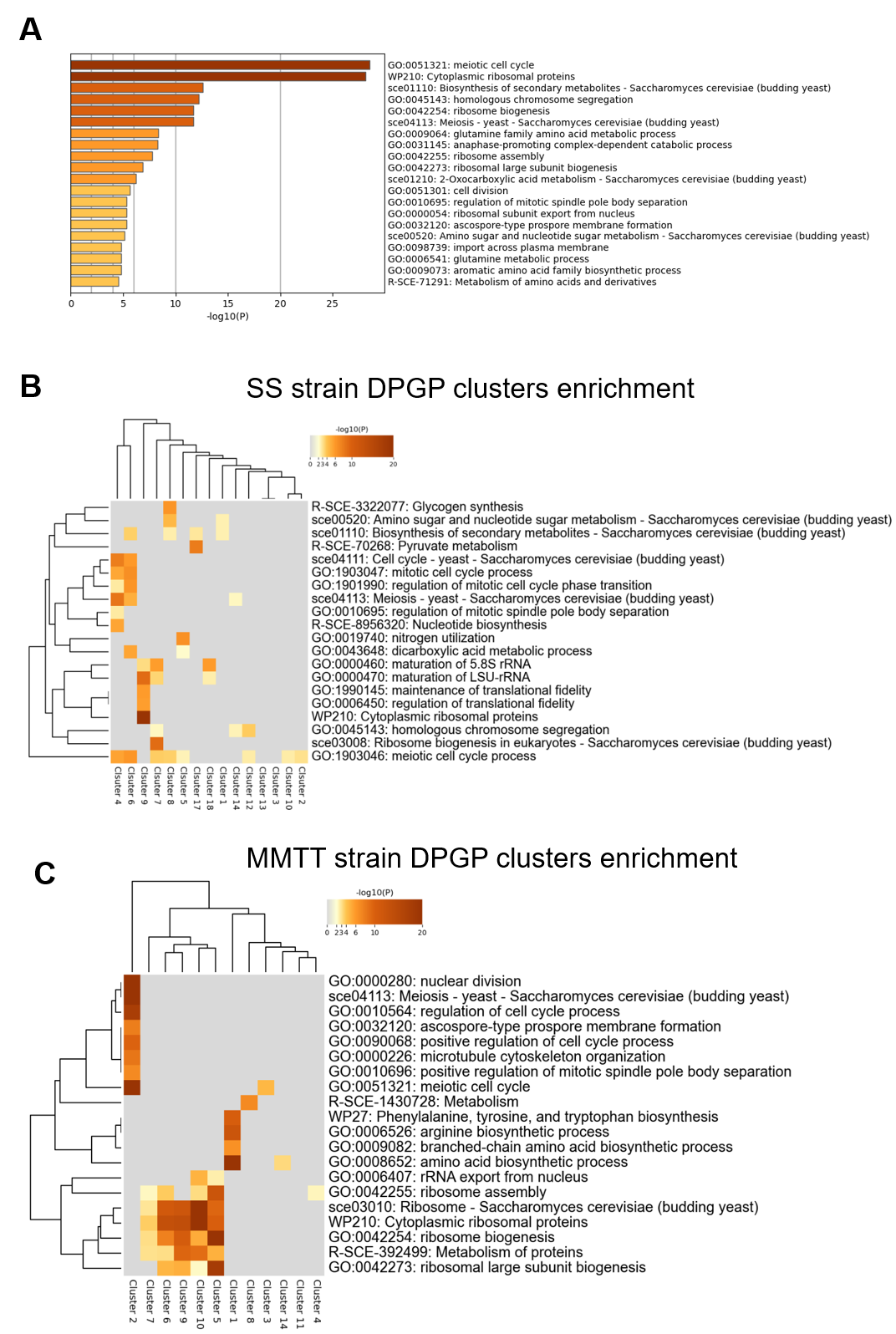

**Supplementary Fig. 4: Metascape enrichment analysis reveals distinct pathway enrichment patterns between SS and MMTT strains over time.** (A) GO enrichment analysis was performed on 1,080 differentially expressed genes between SS and MMTT strains, identified using the DESeq2 LRT method. (B-C) Heatmaps display the p-values of enriched GO terms, with hierarchical clustering applied to both rows (GO terms) and columns (clusters obtained through DPGP clustering) for SS and MMTT strains, respectively.

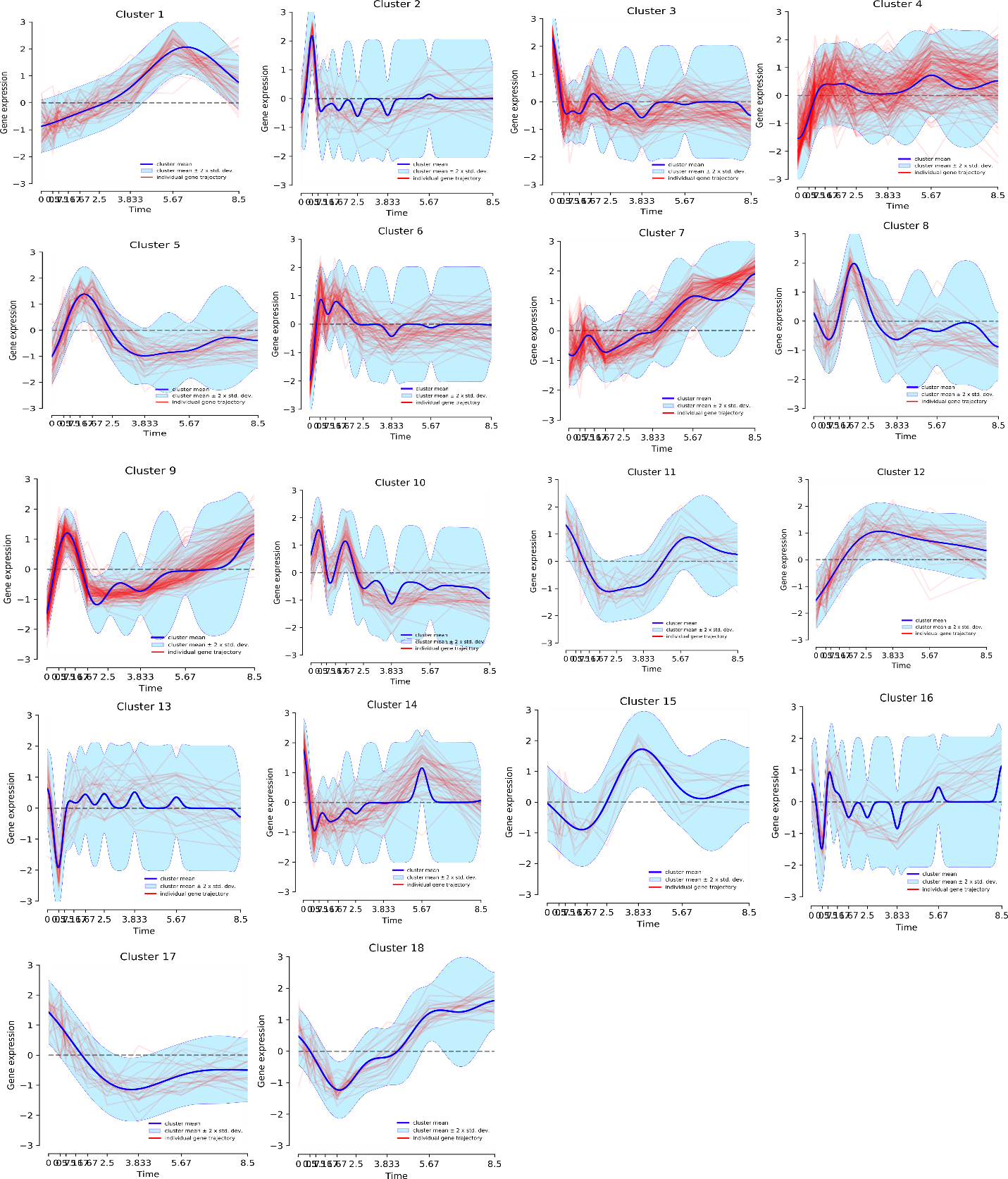

**Supplementary Fig. 5:** (A) Temporal gene expression trajectories of 18 clusters of SS train obtained through DPGP clustering on gene expression profiles of DEGs obtained from the DESeq2 LRT method.

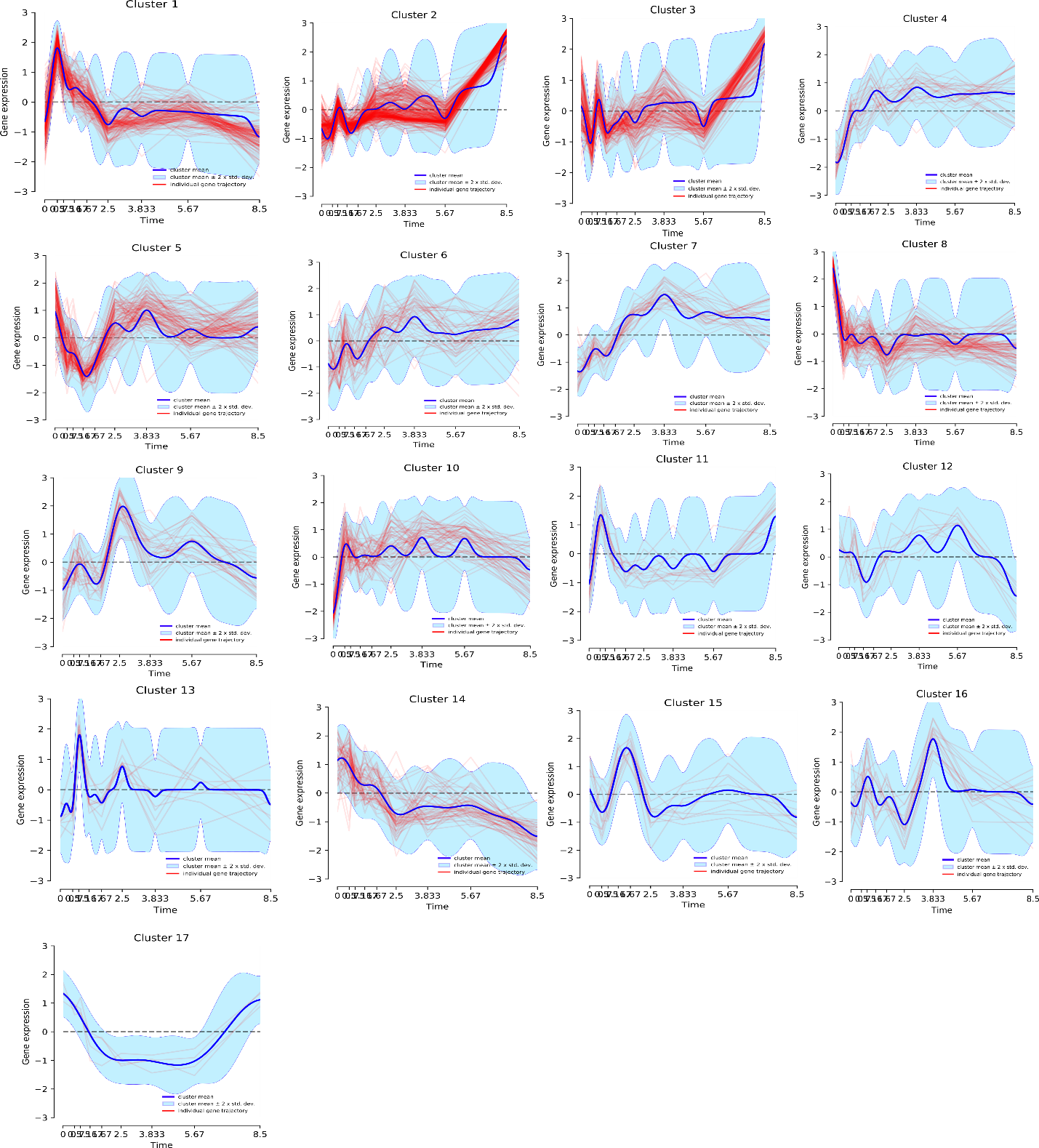

**Supplementary Fig.5** (B)Temporal gene expression trajectories of 17 clusters of MMTT train obtained through DPGP clustering on gene expression profiles of DEGs obtained from the DESeq2 LRT method.

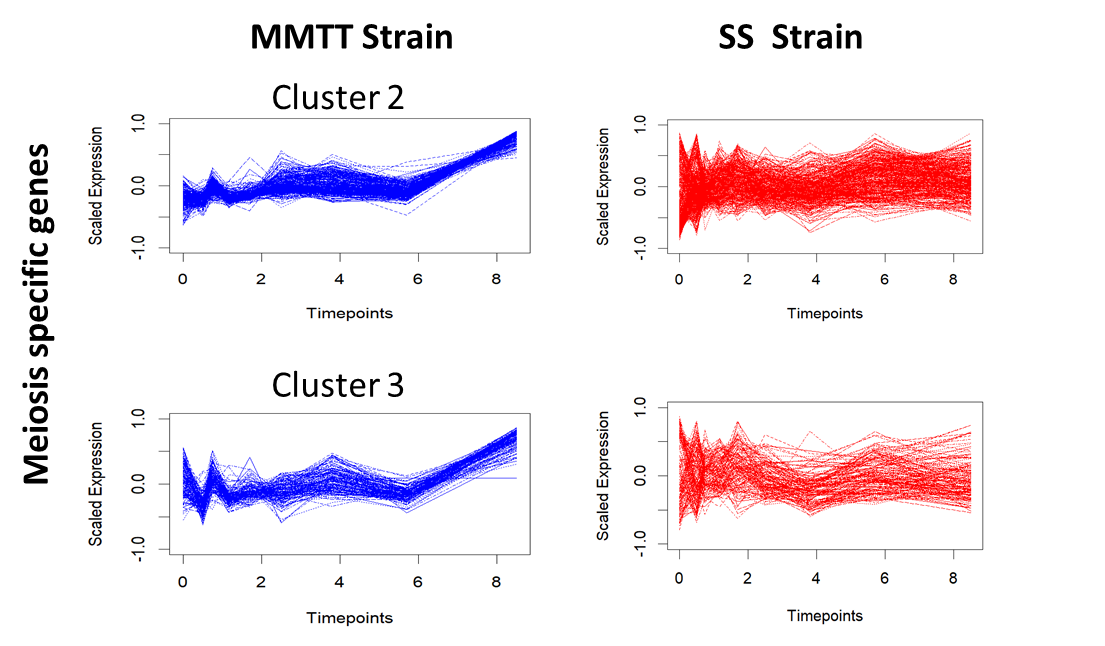

**Supplementary Fig. 6:** Temporal gene expression trajectories of genes in Clusters 2 and 3 of the MMTT train enriched for meiosis-specific pathways and their trajectories in the SS strain.

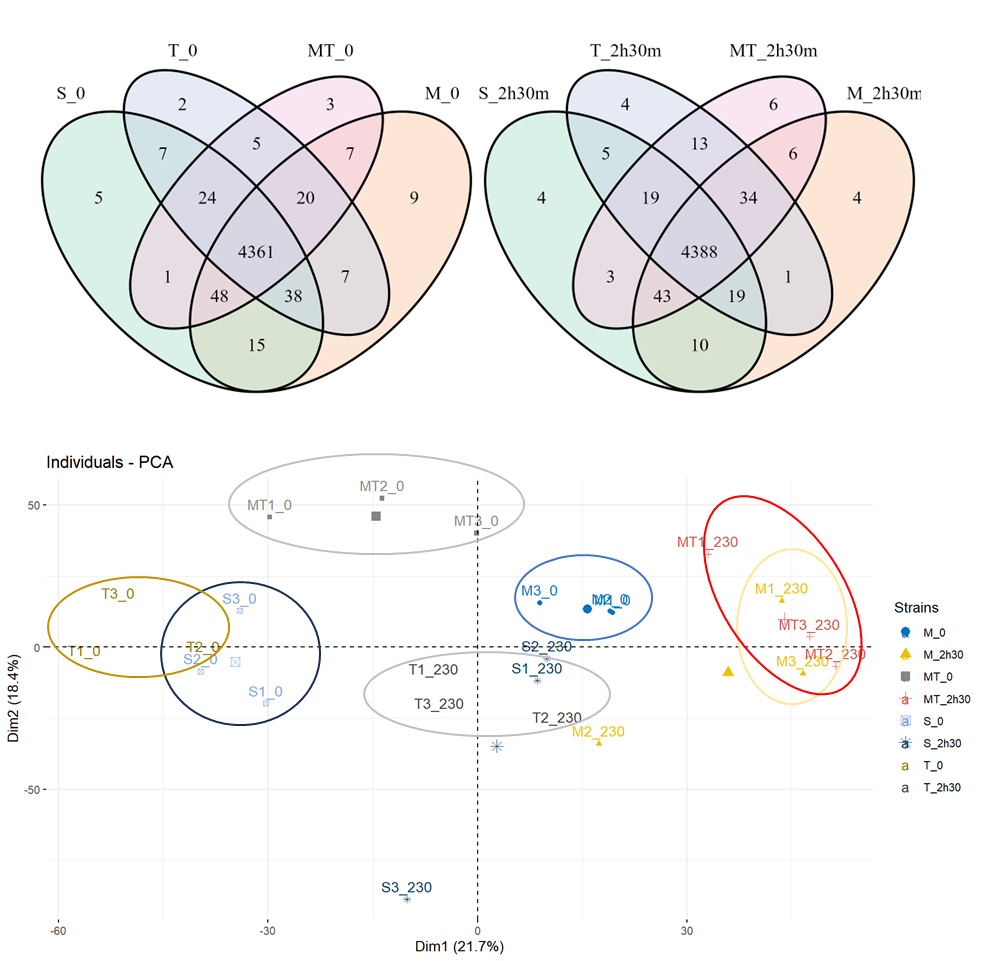

**Supplementary Fig. 7:** PCA of the protein abundance data of SS, MM, TT and MMTT strains during the 0 h and 2 h 30 min into sporulation. The analysis identified one of the S288c strain replicates at 2 h 30 min as a potential outlier, which was excluded from further analysis.

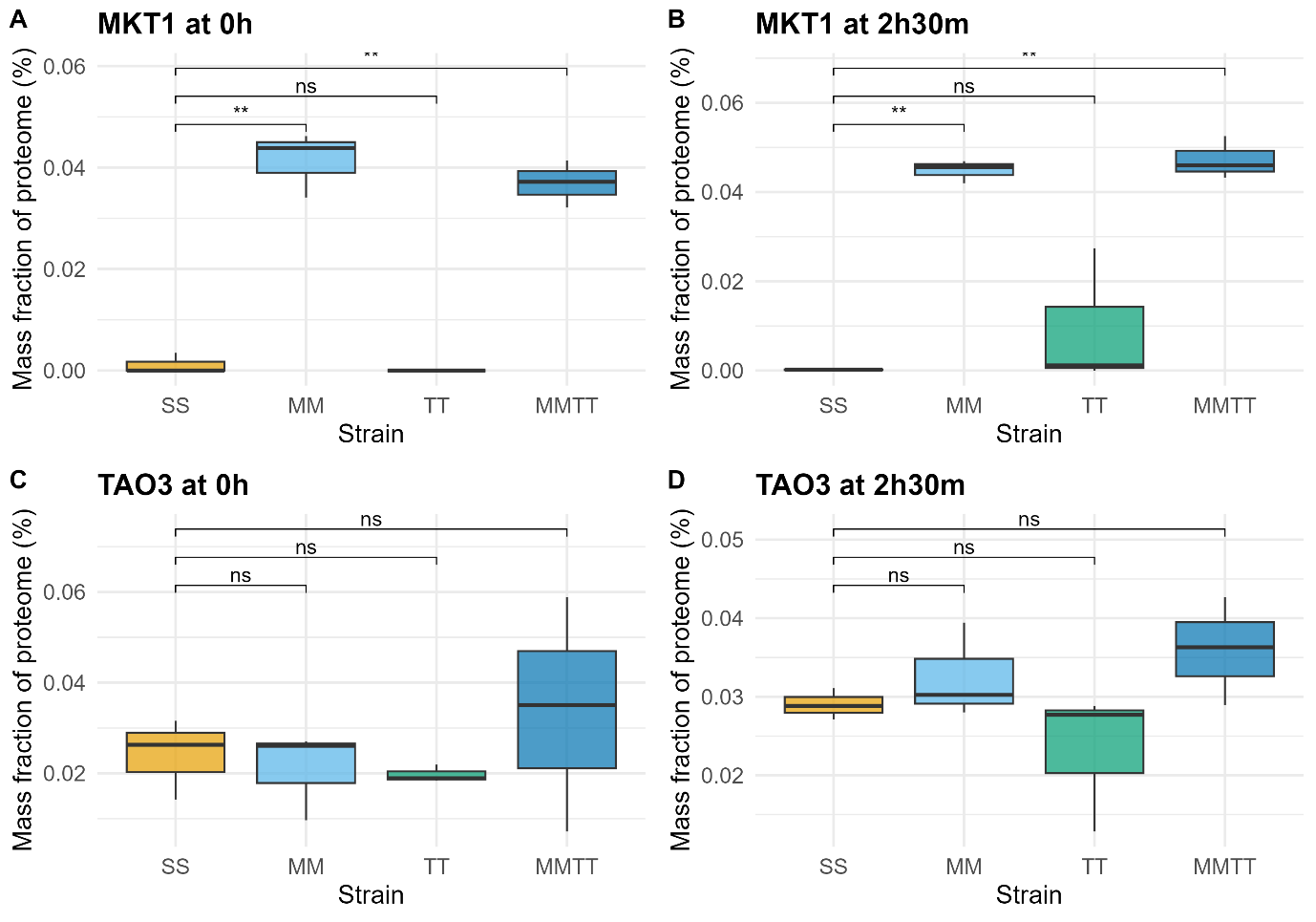

**Supplementary Fig. 8:** **Protein expression levels of MKT1 and TAO3 across yeast strains and time points.** Boxplots showing the mass fraction of the proteome (%) allocated to the *MKT1* (panels A, B) and *TAO3* (panels C, D) and in four yeast strains (SS, MM, TT, and MMTT) at two time points: 0 h (A, C) and 2 h 30 min (B, D) after sporulation induction. Statistical comparisons were performed using unpaired t-test between SS and each of the other strains (MM, TT, MMTT). Significance levels are indicated as follows: ns, not significant; *, p < 0.05; **, p < 0.01.

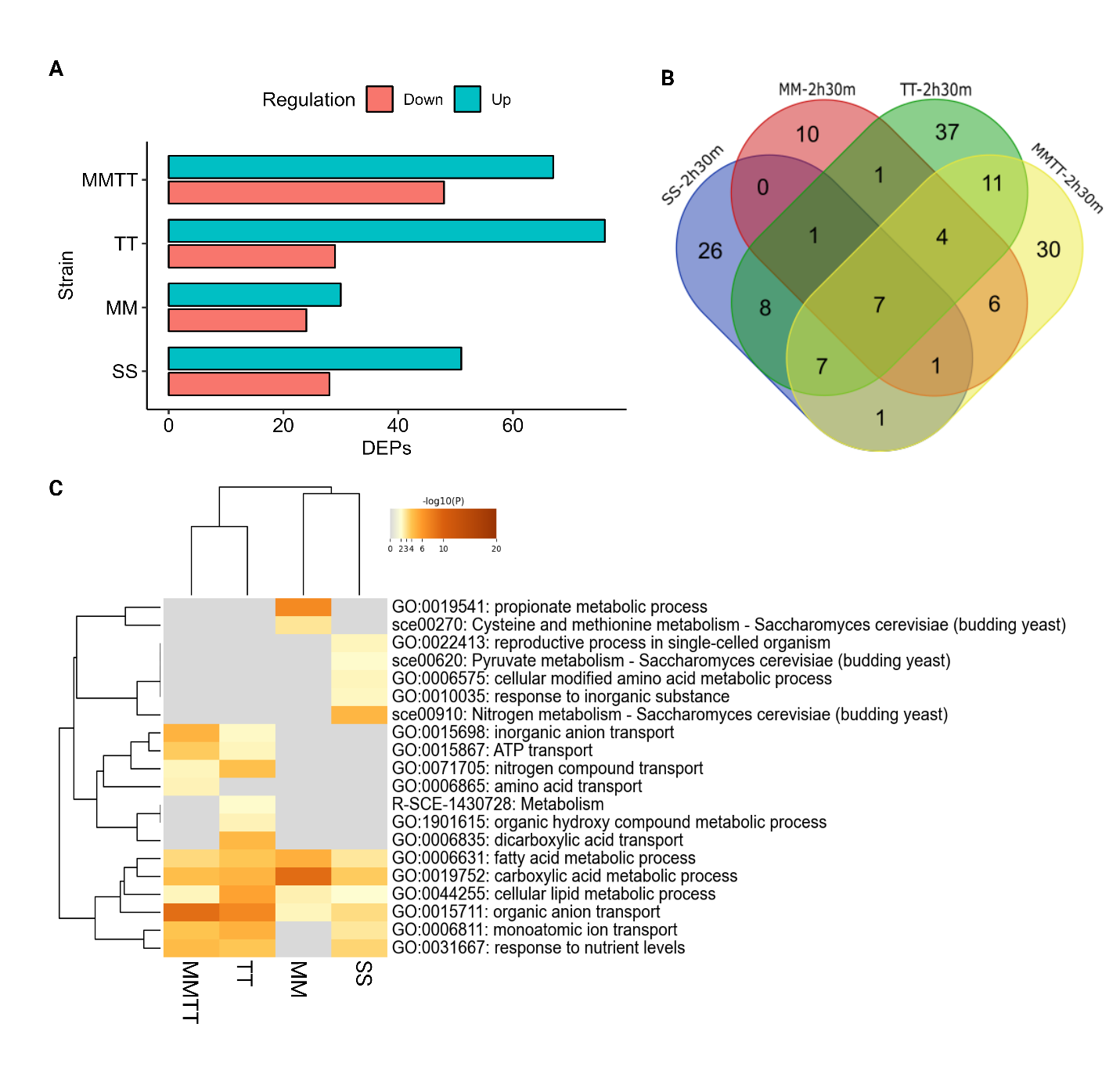

**Supplementary Fig. 9:** **Differentially expressed proteins in the SS, MM, TT and MMTT strains during the early sporulation phase (2 h 30 min).** (A) Bar plots showing the number of significantly differentially expressed genes compared to the initial time point. (B) Venn diagrams depicting the number of proteins specifically upregulated and down-regulated identified through comparison of the differentially expressed proteins in the three phases of yeast growth. (C) Enrichment GO-terms for upregulated proteins in SS, MM, TT and MMTT strains during 2 h 30 min in comparison with the 0 h.

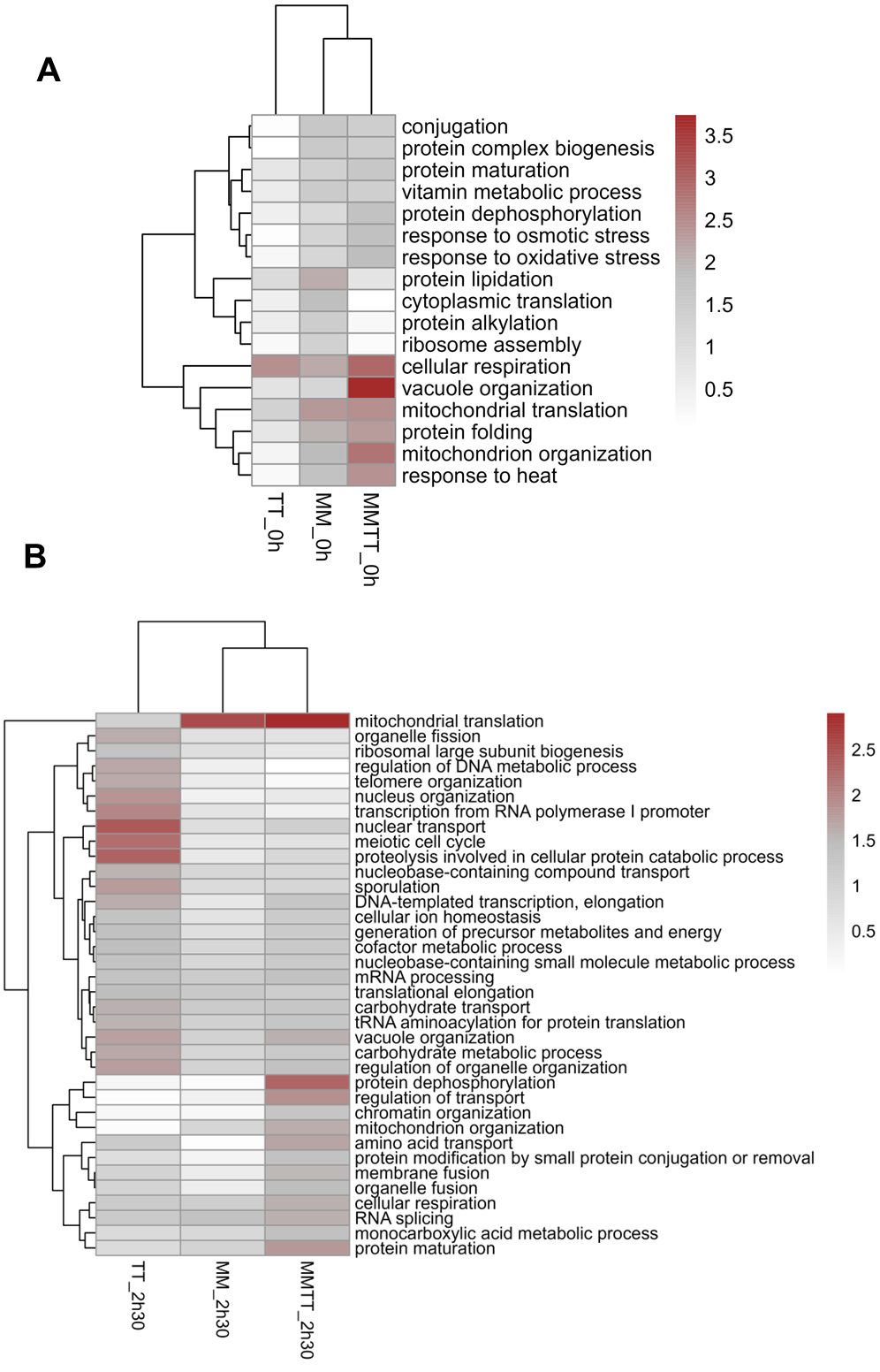

**Supplementary Fig. 10:** Pairwise comparison of protein allocation between SS and allele replacement strains (MM, TT, and MMTT) for key GO-mapper terms at (A) 0 h and (B) 2 h 30 min. GO terms with significant differences in at least one pairwise comparison are shown.

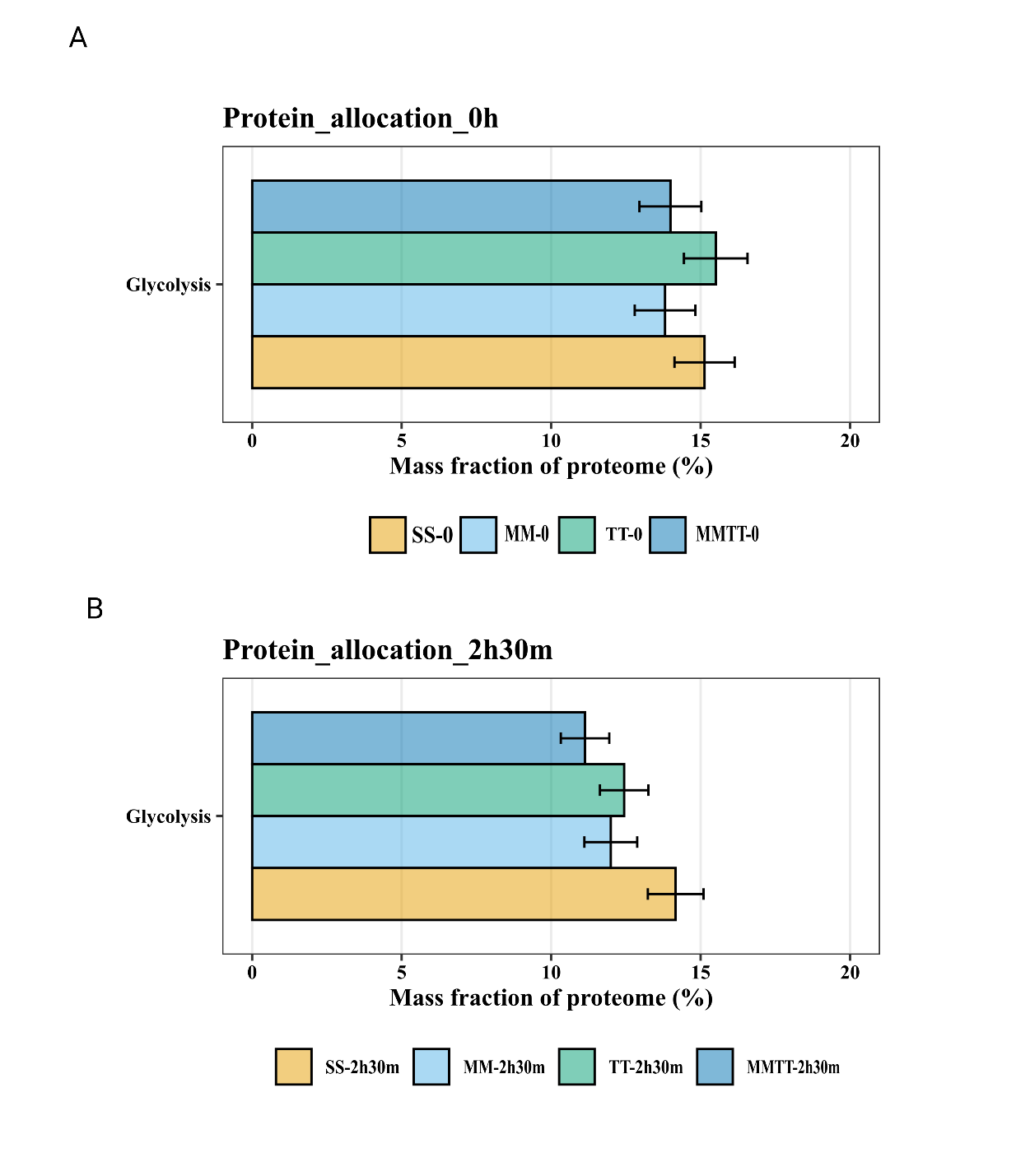

**Supplementary Fig. 11:** **Protein allocation to glycolysis decreases during sporulation in the MMTT strain.** Allocation of the whole cellular proteome to glycolysis pathway in SS, MM, TT and MMTT during the (A) 0 h and (B) early stage of sporulation (2 h 30 min), calculated as the mean percentage allocation. Data are mean ± SD of three biological replicates (except the S strain at 2 h 30 min, which had two replicates).

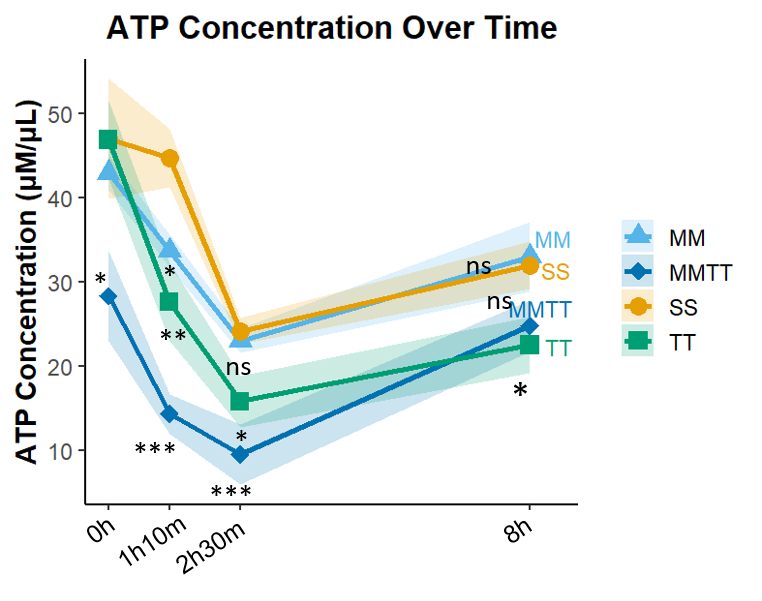

**Supplementary Fig. 12: Intracellular ATP concentrations over time in sporulation medium.** ATP levels were quantified in strains SS, MM, TT and MMTT at 0 h, 1.16 h, 2.5 h, and 8 h. Data are presented as mean ± SD from 2 to 3 biological replicates. Statistical significance was assessed using unpaired two-tailed t-tests for M, T and MT in comparison with the S strain (*p < 0.05, **p < 0.01, ***p < 0.001; ns, not significant) (n = 2 or 3).

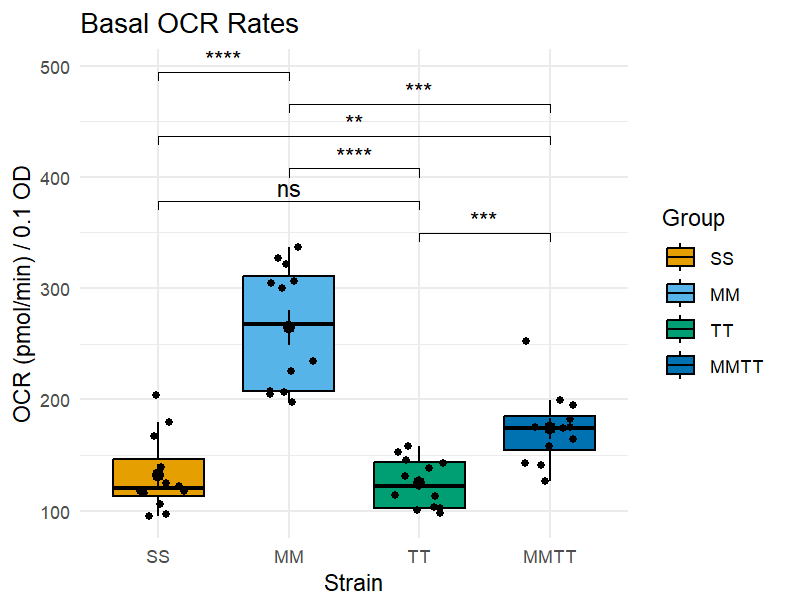

**Supplementary Fig. 13**: **Basal oxygen consumption rate (OCR) across yeast strains.** Boxplots show the normalised basal OCR (pmol/min per 0.1 OD) for each strain: SS, MM, TT, and MMTT. Each point represents an individual three Seahorse measurement under acetate medium conditions. Statistical comparisons were performed using pairwise t-tests with Bonferroni correction. Significant differences between groups are marked with asterisks (* p < 0.05, ** p < 0.01, *** p < 0.001, **** p < 0.0001).

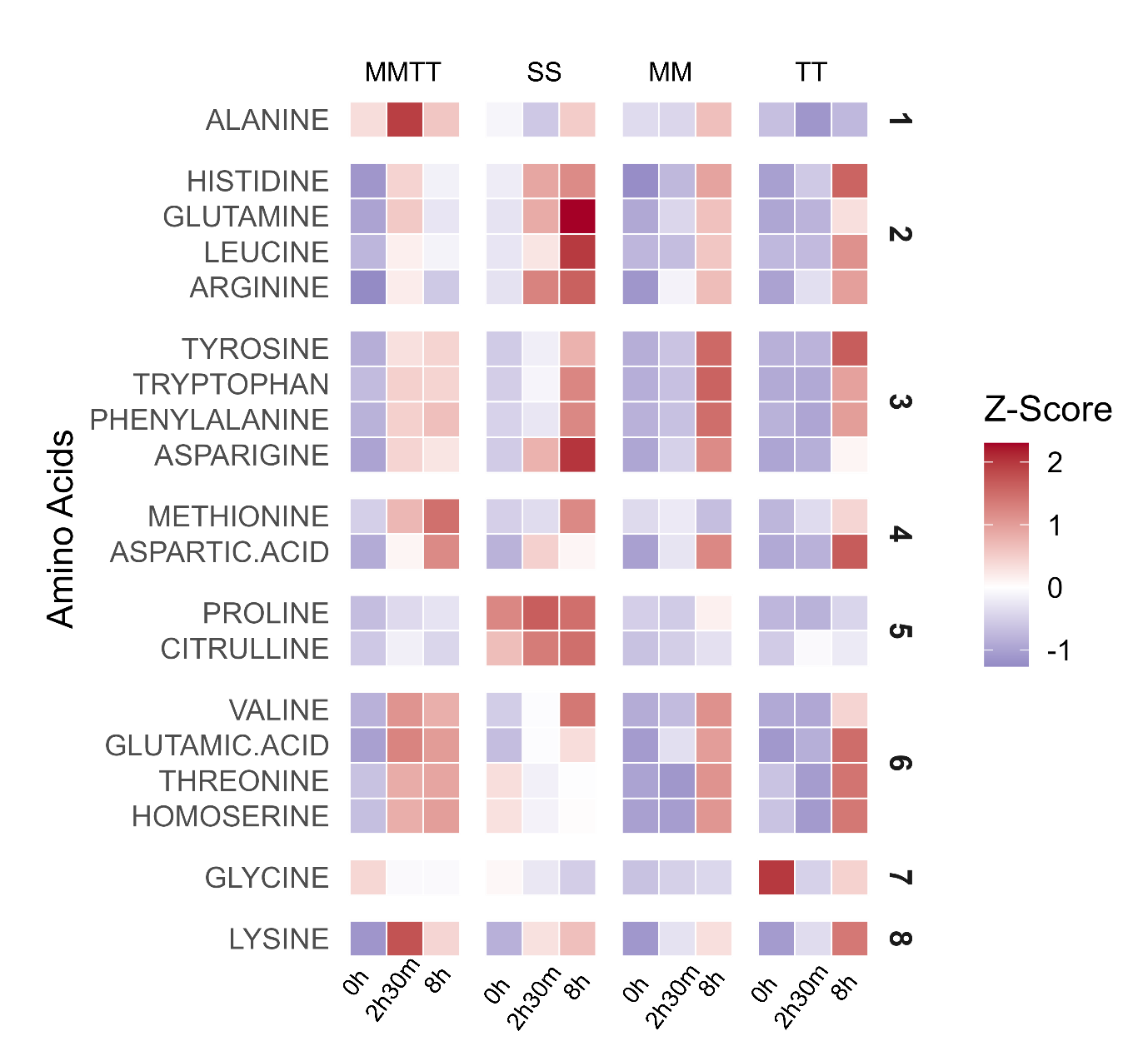

**Supplementary Fig. 14:** Heatmap showing the z-scores of amino acid intensity values across time points and strains. Amino acids are ordered based on the hierarchical clustering of values in the MMTT strain, keeping time constant.

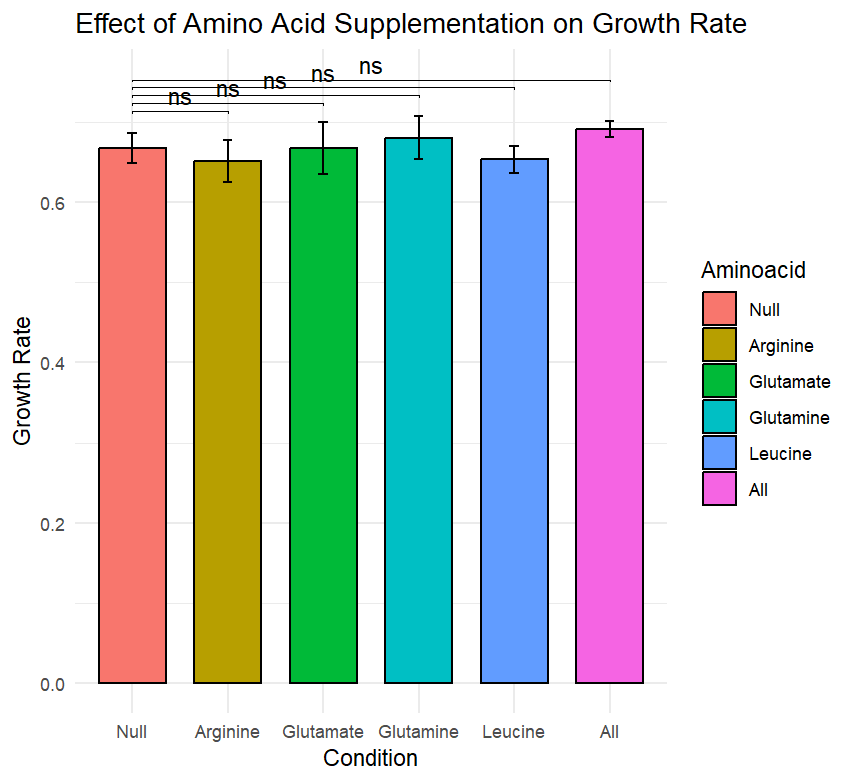

**Supplementary Fig. 15: Amino acid supplementation does not rescue the growth defect of MMTT-*arg4*Δ.** Bar plot showing the growth rates of the MMTT-*arg4*Δ strain in YPD and YPD supplemented with individual amino acids. Growth rates were measured under each condition, and the mean ± SD is shown. Statistical significance was assessed using an unpaired two-sided t-test comparing each supplemented condition to the YPD control. Significance levels are indicated as follows: ns, not significant; *, p < 0.05; **, p < 0.01.

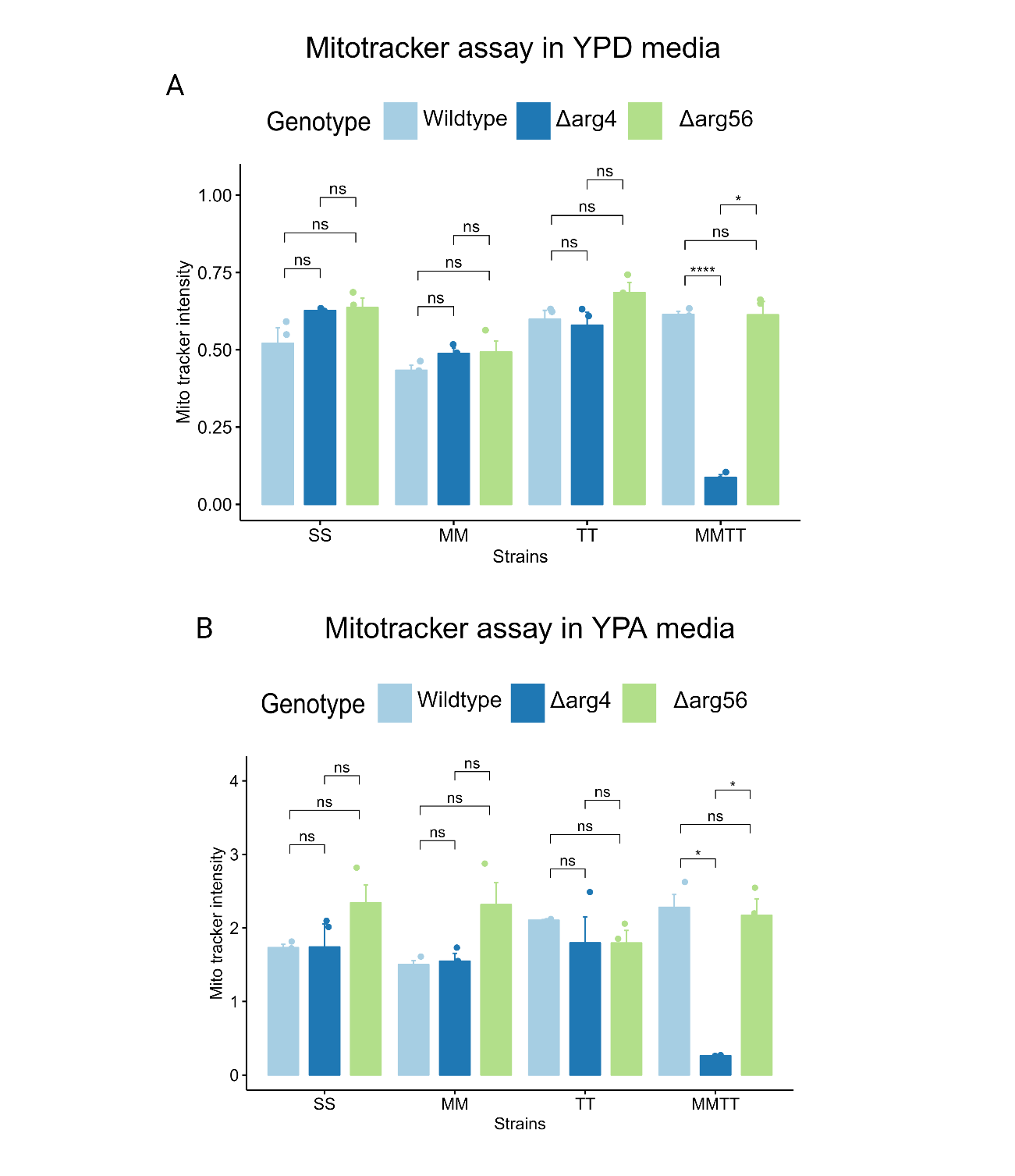

**Supplementary Fig. 16:** Mitotracker fluorescent assay on haploid S, M, T and MT and their respective *arg4* and *arg56* deletions after 2 h incubation in (A) YPD medium and (B) YPA medium. P-values were calculated using an unpaired t-test in the rstatix R package (v.0.7.2). Significance levels are indicated as follows: **** p < 1e^-4^, *** p < 1e^-3^, ** p < 1e^-2^, * p < 5e^-2^, and ns (not significant) for p > 5e^-2^. The mean ± S.E. of at least three biological replicates is shown.

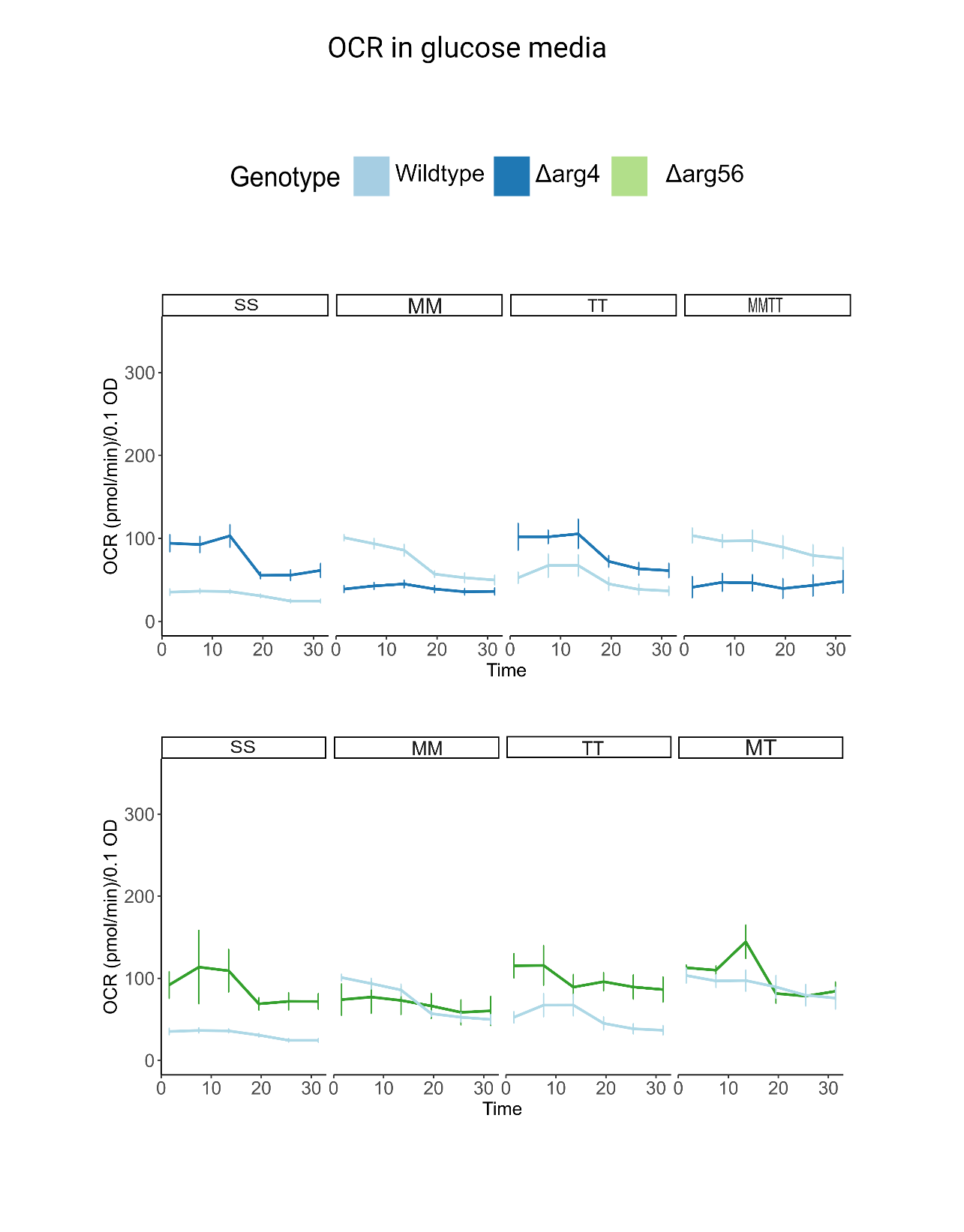

**Supplementary Fig. 17:** The oxygen consumption rate (OCR) of SS, MM, TT and MMTT for wildtype (n = 4), *arg4Δ* (n = 4) and *arg56Δ* (n = 3) cells was measured using the Seahorse Extracellular Flux 96 Analyzer, grown in glucose media. After three basal measurements, sodium azide was injected into the wells to shut off mitochondrial oxygen consumption, and an additional three sets of measurements were taken.

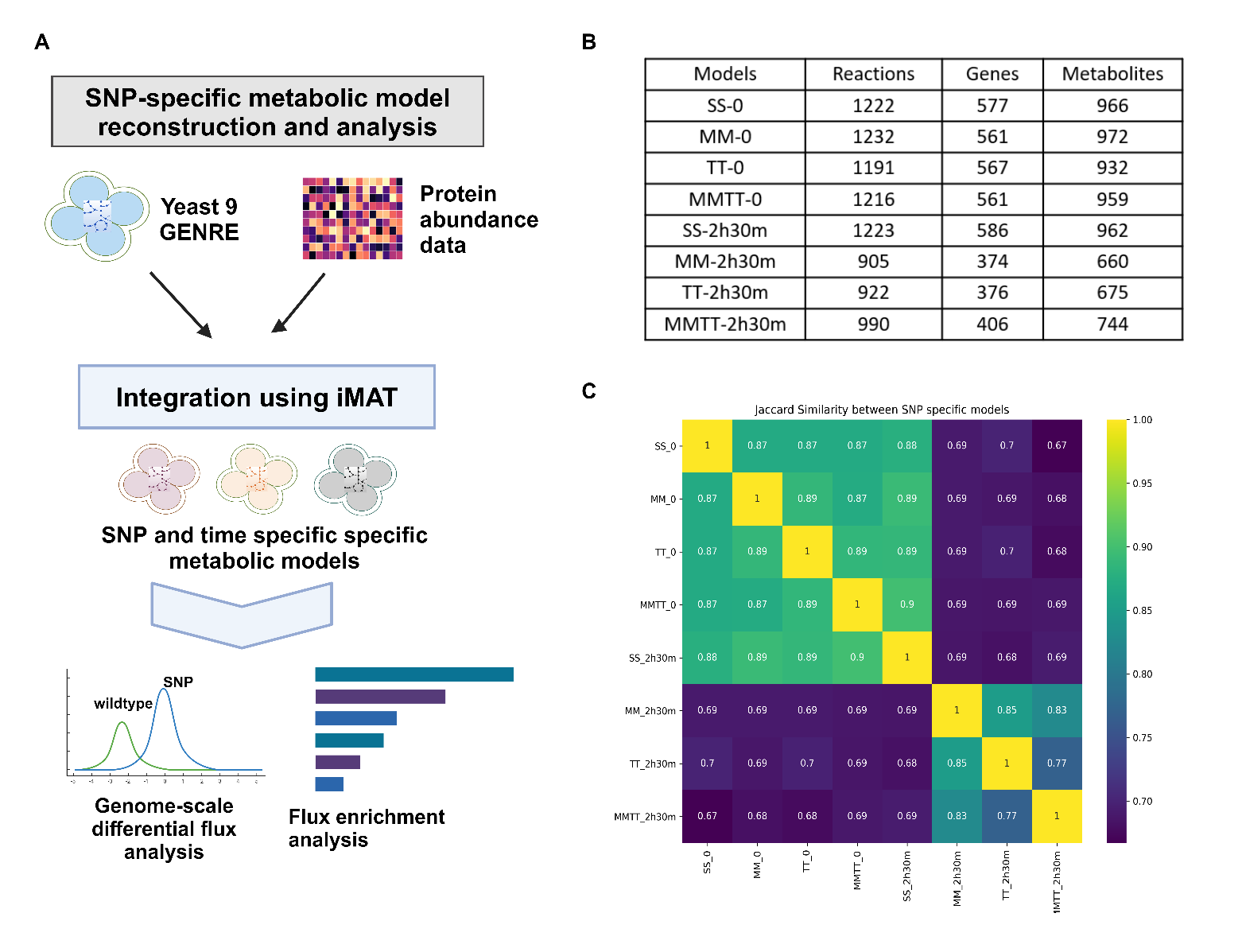

**Supplementary Fig. 18**: Context-specific models show metabolic heterogeneity during sporulation. (A) Schematic representation of steps involved in generating and analysing SNP and time-specific models. (B) The number of reactions, genes and metabolites present in each context-specific model. (C) Jaccard similarity index of context-specific models based on the presence or absence of reactions.

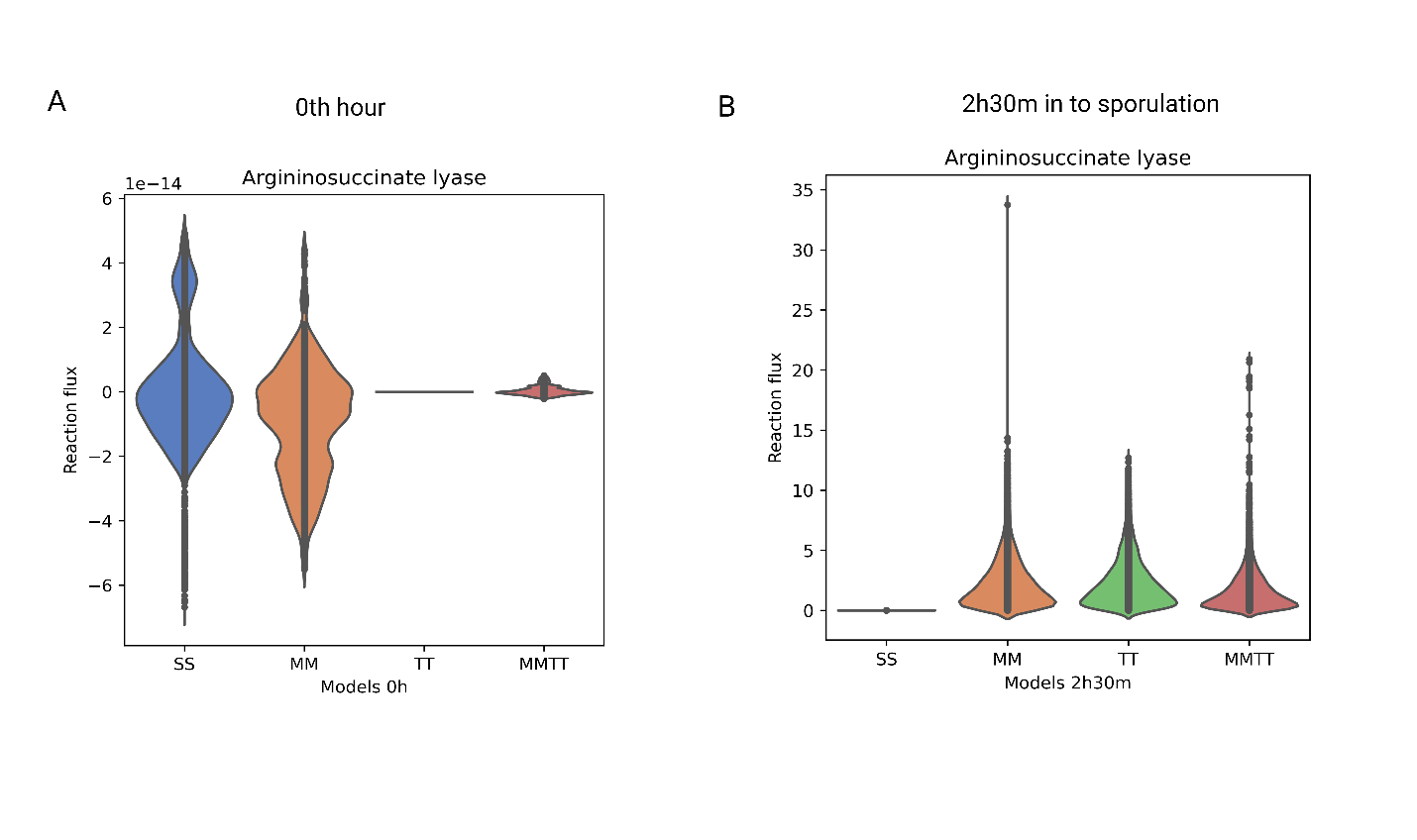

**Supplementary Fig. 19.** The violin plots represent the flux distribution obtained by optGpSampler for Arginosuccinate lyase (*ARG4*) reaction during the (A) 0 h and (B) 2 h 30 min into sporulation in SS, MM, TT and MMTT models.

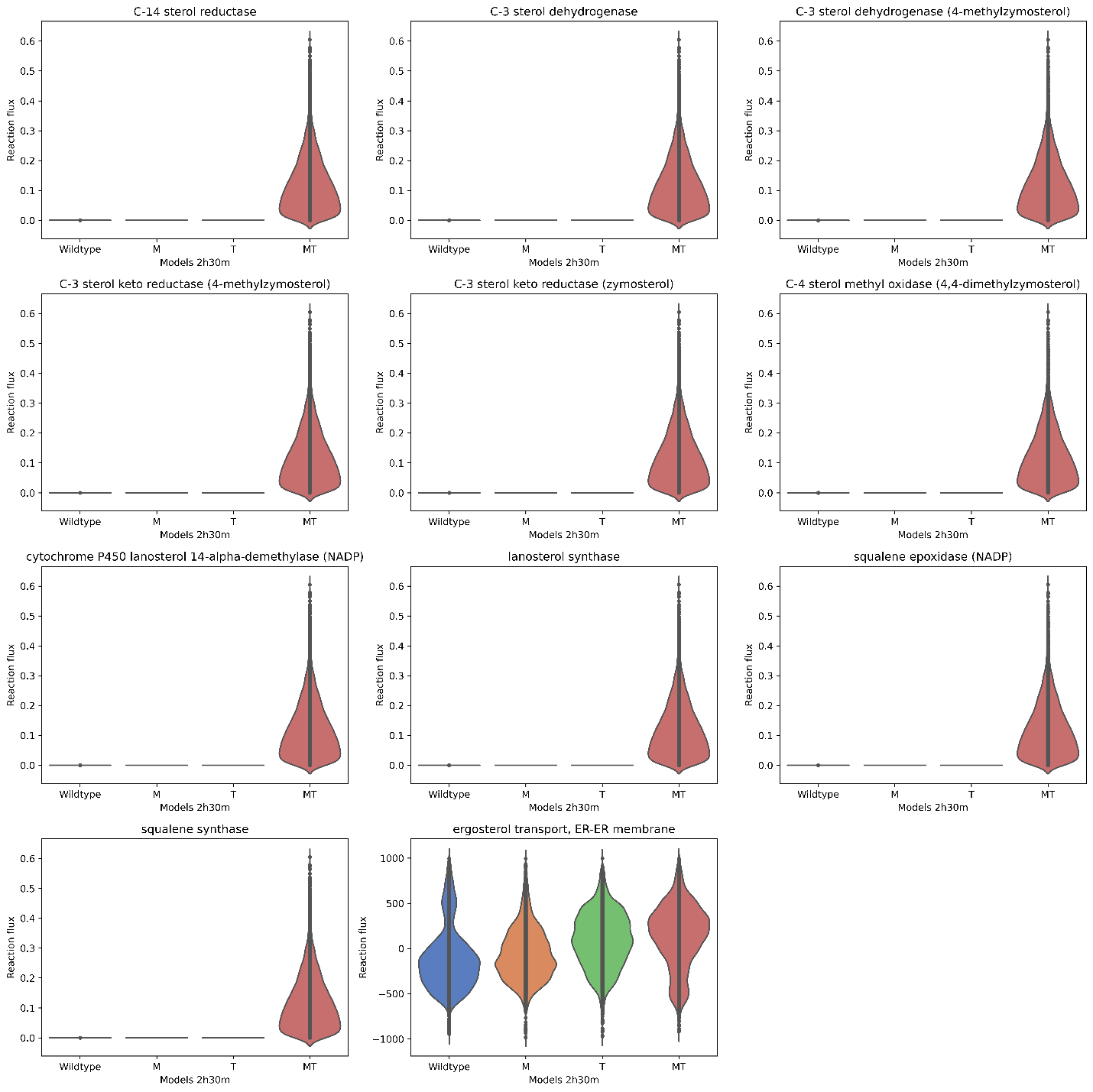

**Supplementary Fig. 20.** Steroid biosynthesis pathway is differentially activated when *MKT1^89G^* and *TAO3*^4477C^ interact. The violin plots represent the flux distribution obtained by optGpSampler for key reactions in the steroid biosynthesis pathway during 2 h 30 min into sporulation in SS, MM, TT and MMTT models.
